# Lung injury epigenetically primes mesenchyme for amplified activation upon re-injury

**DOI:** 10.1101/2022.03.04.483065

**Authors:** Dakota L. Jones, Nunzia Caporarello, Jeffrey A. Meridew, Kyoung M. Choi, Andrew J. Haak, Patrick A. Link, Qi Tan, Tamas Ordog, Giovanni Ligresti, Daniel J. Tschumperlin

## Abstract

The lungs have a remarkable capacity to repair. However, repetitive injury can lead to progressive fibrosis and end-stage organ failure. Whether tissue-resident mesenchymal cell populations retain epigenetic memory of prior injuries that contribute to this pathological process is unknown. Here we used a genetic lineage labeling approach to mark the lung mesenchyme prior to injury, then performed multi-modal analyses on isolated lung mesenchyme during the initiation, progression and resolution of the fibrotic response. Our results demonstrate the remarkable epigenetic and transcriptional plasticity of the lung mesenchyme during fibrogenic activation and de-activation. Despite this plasticity, we also find that the lung mesenchyme retains specific epigenetic traits (memory) of prior activation, resulting in amplified induction of a fibrogenic program upon re-injury. We identify Runx1 as a critical driver of both fibrogenic activation and epigenetic memory. Comparison of fresh isolated and cultured lung mesenchyme demonstrates that Runx1 is spontaneously activated in standard culture conditions, previously masking these roles of Runx1. Genetic and pharmacological targeting of Runx1 dampens fibrogenic mesenchymal cell activation in cell and tissue models, confirming its functional importance. Finally, publicly available scRNAseq data reveal selective expression of Runx1 in the fibrogenic cell subpopulations that emerge in mouse and human fibrotic lung tissue. Collectively, our findings implicate Runx1 in both the initiation and memory of fibrogenic mesenchymal cell activation that together prime amplified mesenchymal cell responses upon repeated injury.

## Introduction

The lung presents the largest surface area of the body exposed to pathogens, particles and noxious gases. Following injury, repair or regeneration of the alveolar gas exchange regions of the lung is required to sustain lung function. Alveolar regeneration requires activation of tissue-resident progenitor cell populations to orchestrate repair (Basil *et al.*, 2020). The lung mesenchyme plays key roles by coordinating extra-cellular matrix repair, while also providing critical paracrine signals within the alveolar niche (Lee *et al.*, 2017; Zepp *et al.*, 2017). In chronic human interstitial lung diseases (ILDs), such as idiopathic pulmonary fibrosis (IPF), mesenchymal cells are locked in persistently activated states that result in distortion and destruction of the alveolar architecture, leading to scarring and progressive decline in organ function (Martinez *et al.*, 2017). Understanding how the lung mesenchyme arrives at this aberrant state of persistent activation is critical to developing more effective therapies and early interventions that promote functional repair.

The etiology of IPF remains poorly understood, but is thought to represent an end-stage result of repeated micro-injuries over time, eventually leading to progenitor cell exhaustion and emergence of aberrant cell types including fibrogenic and contractile fibroblasts in the distal lung (Martinez *et al.*, 2017). In lung fibrosis these activated fibroblasts play multiple pathogenic roles, including the building of the dense extracellular matrix (ECM) scar which limits gas-exchange function, as well as paracrine signaling that promotes aberrant cellular states in other cell compartments (Buechler, Fu and Turley, 2021; Kathiriya *et al.*, 2021). Key aspects of lung fibrosis can be studied in mouse models, the most common of which involves the administration of a single intratracheal dose of bleomycin. The resulting lung injury generates a robust but typically transient fibrogenic response in the distal lung which resolves over time (Jun and Lau, 2018; Tan *et al.*, 2021), providing potentially unique insights into reparative cell states and memory of prior injury. Intriguingly, repetitive dosing of bleomycin results in a persistent fibrotic tissue state, more closely mimicking IPF disease pathology (Degryse *et al.*, 2010; Singh *et al.*, 2017; Burman *et al.*, 2018; Elizabeth F Redente *et al.*, 2021). While the responses to, and long term consequences of, single and repetitive lung injury have been well characterized in other compartments of the lung (e.g. epithelial, immune) (Misharin *et al.*, 2017; Elizabeth F Redente *et al.*, 2021), the extent to which single and repetitive injury affects long-term behavior of the lung mesenchyme is not fully understood.

There is both historical and emerging evidence that structural cells accumulate memories of prior injury. Nearly a decade ago it was shown that a portion of hepatic stellate cells that are activated by liver injury undergo deactivation after successful resolution of fibrotic injury but retain an altered state that accelerates subsequent in vitro activation (Kisseleva *et al.*, 2012). More recently, epithelial stem cells of the skin have been shown to retain epigenetic memory of prior tissue injury and inflammation (Naik *et al.*, 2017; Gonzales *et al.*, 2021; Larsen *et al.*, 2021). In both cases, such memory is thought to provide an evolutionary advantage and protective role by accelerating wound repair following a second tissue insult. A potential adverse consequence of such a mechanism is “priming” tissue-resident cells, such as the mesenchyme, for fibrosis if subsequent insults are not successfully resolved or orchestrated. Notably, whereas in vitro studies have demonstrated that mesenchymal cells are capable of retaining memory of previous microenvironments (Balestrini *et al.*, 2012; Yang *et al.*, 2014; Li *et al.*, 2017), in vivo evidence that lung mesenchymal cells acquire and retain a memory of prior injury, or that such memories impact subsequent cellular responses, has been lacking.

Using a time-course multi-modal sequencing approach, we characterized epigenetic and transcriptional features of the lung Col1a2-lineage following a single dose of bleomycin. We found the lung mesenchyme exhibits extensive epigenetic and transcriptional plasticity during fibrotic lung injury, fibrosis and resolution. We further identified Runx1 as a key player in these epigenetic and transcriptional changes. Despite the evident plasticity, we also found evidence that the mesenchyme retains an epigenetic and transcriptionally “poised” state strongly marked by a Runx1 signature, and upon re-injury Runx1 target genes exhibit amplified responses, demonstrating a role for Runx1 in fibrogenic memory. Remarkably, ex vivo culture of lung mesenchyme activates Runx1 expression and Runx1-dependent fibrogenic functions, a phenomenon that has previously obscured the in vivo relevance of Runx1. Finally, using publicly-available mouse and human scRNAseq datasets of fibrotic lung tissue we identify the selective emergence of Runx1 expression in fibrogenically activated cell populations in fibrotic mouse and human lung tissue. Collectively, these results define the remarkable plasticity of the lung mesenchyme as well as the importance of fibrogenic mesenchymal memory in amplified responses to repeated injury.

## Results

### Transient expansion and activation of the Col1a2-lineage following a single intratracheal administration of bleomycin

To label and trace the collagen I expressing cells in the lung we utilized an inducible Col1a2-CreERT:Rosa26-mTmG mouse which, following tamoxifen administration, fluorescently labels Col1a2+ cells and their progeny (Fig. S1).

To study changes in the abundance and genomic landscape of the Col1a2-lineage during lung fibrogenesis and resolution, we induced recombination and two weeks later injured the lungs by intratracheal administration of bleomycin (Fig. 1a, b). Flow-cytometry analyses revealed a ~50% increase in the relative abundance of the Col1a2-lineage population at 14 days post injury (dpi), which returned to baseline levels at 28 and 56 dpi (Fig. 1c), consistent with prior reports of myofibroblast cell clearance that accompanies fibrosis resolution (Elizabeth F. Redente *et al.*, 2021). Analysis of transcripts involved in cell-cycle programs in FACS isolated Col1a2-lineage mesenchyme confirmed robust but transient increases at 14 dpi, consistent with cytometry results (Fig. 1d). These data confirm that robust proliferation of the lung resident Col1a2-lineage is an early event following fibrotic lung injury. Quantification of lung hydroxyproline content as an indicator of tissue fibrosis revealed a sharp increase at 14 dpi that was maintained at 28 dpi, but then returned to near baseline levels by 56 dpi (Fig. 1e), consistent with other studies showing spontaneous resolution of one time bleomycin injury in young mice (Strunz *et al.*, 2020; Elizabeth F Redente *et al.*, 2021; Tan *et al.*, 2021). Analysis of ECM-related transcripts was consistent with the hydroxyproline data and demonstrated robust but transient elevations in ECM transcript levels in the lung resident Col1a2-lineage post-bleomycin induced injury (Fig. 1f).

**Figure 1:**
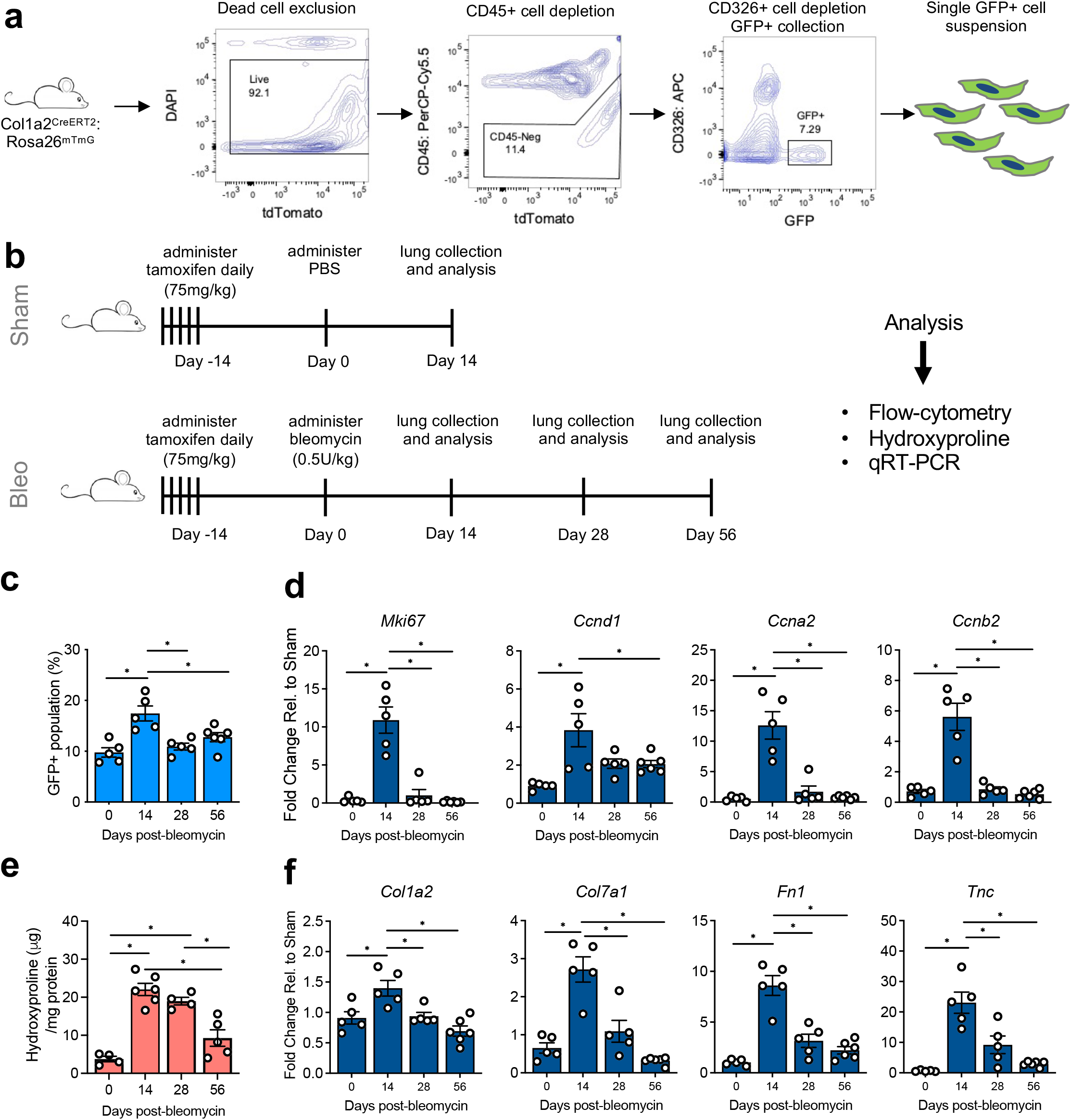
Transient expansion and activation of the lung Col1a2-lineage following a single administration of bleomycin. (a) FACS gating strategy to selectively isolate lineage labeled Col1a2 cells. (b) Experimental approach to study response of the Col1a2-lineage over time following intratracheal administration of bleomycin. (c) Flow-cytometry analysis of the lineage-labeled Col1a2 GFP+ population following bleomycin administration. Data represented as percent of total cell population following CD45+ cell depletion. (d) qRT-PCR analysis of proliferation associated genes from FACS isolated Col1a2-lineage cells following bleomycin. (e) Hydroxyproline analysis of lung tissue following bleomycin. (f) qRT-PCR analysis of ECM associated genes from FACS isolated Col1a2-lineage cells following bleomycin. Each dot represents data obtained from a single mouse (biological replicate). Data represent mean +/- SEM. *P<0.05 evaluated by one-way ANOVA with Tukey’s correction for multiple comparison test.

### ATACseq reveals dynamic and largely reversible chromatin alterations in the Col1a2-lineage after lung injury

Having confirmed that the Col1a2-lineage transiently expands and elevates ECM gene expression after bleomycin injury, we sought to globally define the accompanying chromatin landscape and transcriptional changes by performing ATACseq and RNAseq at 14 and 56 dpi (Fig. 2a). ATACseq analyses of the Col1a2-lineage at 14 dpi compared to sham revealed 17,333 genomic loci increased accessibility whereas 8,185 sites decreased accessibility (Fig. 2b,c). De novo transcription factor (TF) motif enrichment analysis of the genomic loci exhibiting increased accessibility at 14 dpi implicated candidate transcription factors Fosl1, Runx1, and Egr2, consistent with prior reports implicating these factors in tissue fibrosis (Fig. 2d) (Fang *et al.*, 2011; Wernig *et al.*, 2017; Koth *et al.*, 2020; O’Hare *et al.*, 2021). Motif analysis in the genomic regions exhibiting decreased accessibility at 14 dpi implicated loss of candidate transcription factors Twist2, Nfix, and Creb5 (Fig. 2d). Transcriptomic analyses at 14 dpi revealed increased transcripts from 1,589 genes and decreased transcripts from 1,296 genes (Fig. S2a). Pathway analyses of these two gene sets confirmed an increase in cell-cycle and fibrosis related gene programs (Fig. S2b, e), aligned with our FACS and hydroxyproline analyses (Fig. 1c, e), in addition to a decrease in CREB signaling (Fig. S2c, e), consistent with motif enrichment from the ATACseq data (Fig. 2d).

**Figure 2:**
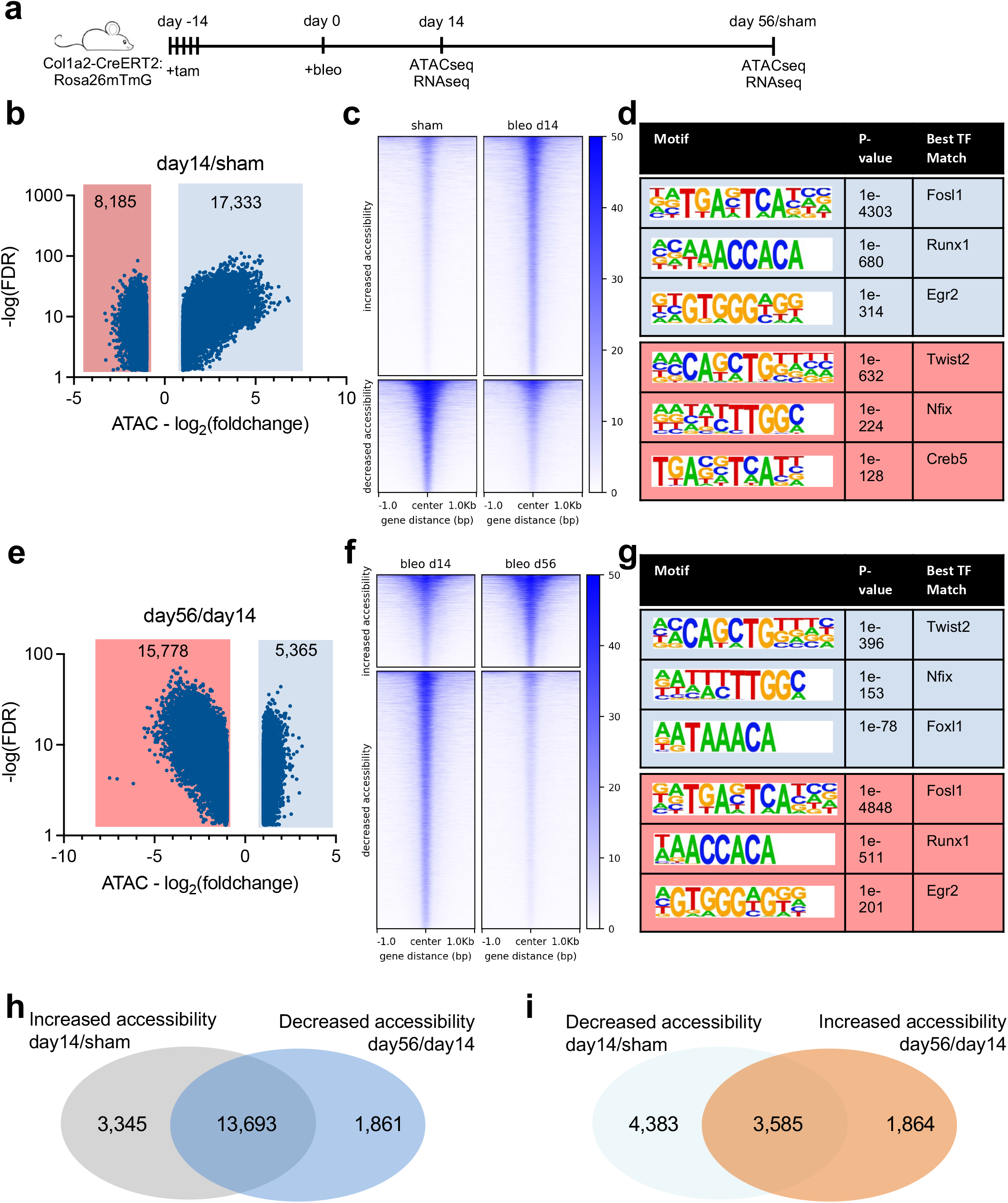
Dynamic chromatin and transcriptional changes accompany Col1a2-lineage activation and deactivation. (a) Experimental approach for multi-modal time course analysis of Col1a2-lineage activation and deactivation following bleomycin administration. (b) Volcano plot of differential accessibility sites of the Col1a2-lineage at 14 dpi compared to sham (n=4 biological replicates/condition). Blue dots represent genes which met differential expression criteria (FDR ≤ 0.05 and log2(foldchange) ≤ −1 or ≥ 1). (c) Heatmap and intensity profiles of differential chromatin accessibility regions at 14 dpi compared to sham. Heatmap scale represents relative reads per genomic region. (d) De novo transcription factor motif enrichment analysis from the genomic loci which increased and decreased accessibility at 14 dpi compared to sham. Motifs ranked by p-value. (e) Volcano plot of differential accessibility sites of the Col1a2-lineage at 56 dpi compared to 14 dpi (n=4 biological replicates/condition). Blue dots represent genes which met differential expression criteria (FDR ≤ 0.05 and log2(foldchange) ≤ −1 or ≥ 1). (f) Heatmap and intensity profiles of differential chromatin accessibility regions at 56 dpi compared to 14 dpi. Heatmap scale represents relative reads per genomic region. (g) De novo transcription factor motif enrichment analysis from the genomic loci which increased and decreased accessibility at 56 dpi compared to 14 dpi. Motifs ranked by p-value. (h) Venn diagram showing the number of genomic loci which significantly increased accessibility at 14 dpi (compared to sham) which also significantly decreased accessibility at 56 dpi (compared to 14 dpi). (i) Venn diagram showing the number of genomic loci which significantly decreased accessibility at 14 dpi (compared to sham) which also significantly increased accessibility at 56 dpi (compared to 14 dpi).

ATACseq analysis revealed a dramatic reversal of the changes in Col1a2-lineage chromatin accessibility from 14 to 56 dpi. 15,778 genomic loci exhibited decreased accessibility while 5,365 genomic loci exhibited increased accessibility from 14 to 56 dpi (Fig. 2e), largely mirroring the changes observed between sham and 14 dpi. Motif enrichment analyses confirmed largely opposite trends compared to the previous motif analysis (sham vs 14 dpi, Fig. 2d), consistent with transient transcriptional activation and subsequent de-activation (Fig. 2g). Comparative analysis of ATACseq and RNAseq changes from sham vs 14 dpi and 14 dpi vs 56 dpi confirmed that the majority of chromatin accessibility and transcriptomic changes associated with Col1a2-lineage activation at 14 dpi were reversed at 56 dpi (Fig. 2h, i and Fig. S2d, e). Collectively, these results reveal that Col1a2-lineage activation during lung injury and repair is driven by largely transient and reversible epigenomic and transcriptomic changes.

### Multi-modal analyses reveal Runx1 as a regulator of fibrogenic Col1a2-lineage activation

To narrow the list of candidate factors driving fibrogenic Col1a2-lineage activation in vivo, we performed a multi-modal merged analysis of the ATACseq and RNAseq datasets. Briefly, we identified the differential ATAC sites exhibiting increased accessibility at 14 dpi compared to sham that contained the top ranked DNA TF motifs identified in Figure 2d (Fosl1, Runx1, and Egr2) to generate target ATAC regions for each TF. We then annotated each of these regions to the nearest transcriptional start site (TSS) to generate a target gene list for each TF. Next, we retrieved the expression profile for each of these gene lists from the RNAseq analysis (sham vs 14 dpi). Finally, we selected for differential expression (FDR ≤ 0.05) and ran gene ontology analysis to identify pathways regulated by each TF.

Analysis of the pathways generated for each TF revealed Runx1 as the only TF enriched for fibrotic related pathways (“collagen biosynthetic processes”) implicating a role for Runx1 in fibrogenic Col1a2-lineage activation in the lung following injury (Fig 3a). To test whether ATACseq changes overlapped with Runx1 occupancy, we examined publicly-available Runx1 ChIPseq data from mouse embryonic fibroblasts (GSE90893) (Chronis *et al.*, 2017). This analysis identified a 36% overlap (n=6,129) between the open chromatin sites with increased accessibility at 14 dpi and Runx1 occupancy in mouse embryonic fibroblasts (Fig 3b). Fibrosis relevant ECM and matricellular-specific genomic loci with Runx1 occupancy include Col7a1, Col8a1, Col5a1, Cthrc1, and Tnc (Fig 3c). Interestingly, we also observed increased chromatin accessibility in the Runx1 locus at 14 dpi, and this region is also occupied by Runx1 itself (Fig 3c), consistent with a positive feedback loop of Runx1 regulating its own expression within the Col1a2-lineage once activated following injury. RNAseq transcript profiles for these genes confirmed increased expression at 14 dpi (Fig 3d) demonstrating consistency between increased chromatin accessibility, Runx1 occupancy, and increased transcript levels at 14 dpi in the lung resident Col1a2-lineage.

**Figure 3:**
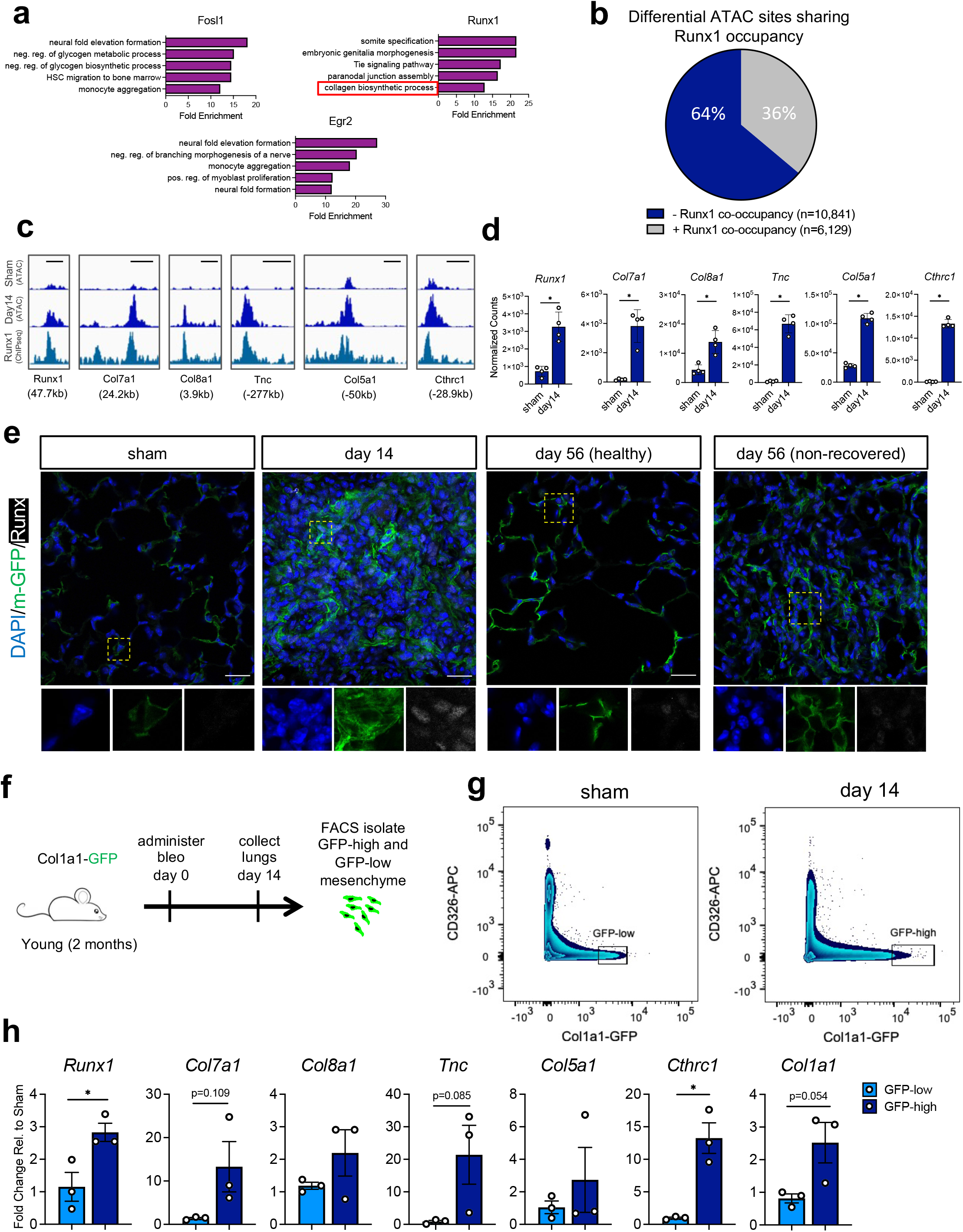
Multi-modal merged analyses reveal Runx1 as a core regulator of Col1a2-lineage cell activation. (a) Overrepresented gene ontology terms from multi-modal analyses for each transcription factor ranked by fold enrichment. (b) Merged occupancy analysis of Runx1 (from previously published ChIPseq (GSE90893)) and increased differential chromatin accessibility regions at 14 dpi compared to sham. (c) Candidate accessibility profiles of the increased differential chromatin accessibility regions at 14 dpi compared to sham and Runx1 occupancy. Scale bars represent 1kb in genomic distance. Numbers below occupancy profiles represents genomic distance to TSS. (d) Expression profiles from RNAseq of genes from panel c. Each dot represents data from a single biological replicate. Error bars represent SEM. *FDR(i.e. P-adj)<0.05 evaluated by Benjamini and Hochberg method. (e) IHC for lineage-traced Col1a2 cells (m-GFP) and Runx1 at 14- and 56-dpi. Yellow dashed boxes demarcate zoomed in regions. Scale bars represent 25 μm. (f) Experimental approach to isolate GFP-high and GFP-low mesenchyme from Col1a1-GFP reporter mice at 14 days following bleomycin administration. (g) Flow cytometry plot showing gating strategy to isolate GFP-high and GFP-low mesenchyme. (h) qRT-PCR analysis of GFP-low and GFP-high lung mesenchyme. Each dot represents data from a single biological replicate. Error bars represent SEM. *P<0.05 evaluated by unpaired t-test.

To assess Runx1 activity in the Col1a2-lineage following bleomycin-induced lung injury, we performed immunostaining of lung sections in sham controls and at 14 and 56 dpi for Runx1. Immunostaining at 14 dpi showed increased levels of nuclear Runx1 in the Col1a2-lineage labeled cells compared to an absence of staining in control mice (Fig. 3e) consistent with increased transcript levels of Runx1 at 14 dpi. At 56 dpi the majority of the lung appeared normal, consistent with broad resolution of fibrosis at 56 dpi (Fig 1e). However, occasional non-fibrotically resolved regions were also observed sporadically throughout the lung at 56 dpi (Fig 3e). Interestingly, immunostaining of nuclear Runx1 was diminished but still apparent in Col1a2-lineage cells in both the healthy and non-resolved regions at 56 dpi. Together, these data complement our ATAC and RNAseq analyses and are consistent with a role for Runx1 in Col1a2-lineage activation at 14 dpi, with a reduced but potentially still active role at 56 dpi.

Runx1 signaling regulates collagen-related mesenchymal cell function in other tissues including those of the musculoskeletal system (Tang *et al.*, 2021). To test whether Runx1 is positively associated with lung mesenchyme that is actively producing collagen, we administered PBS or bleomycin transgenic mice expressing GFP under the control of the Col1a1 promoter (Col1a1-GFP), and sorted for GFP+ lung mesenchyme at 14 dpi (Fig. 3f). As we previously observed (Caporarello *et al.*, 2019, 2020) following bleomycin administration a GFP-high population appeared by FACS analysis that was unique to the injured lung (Fig. 3g). We therefore sorted the highest GFP expressing populations from both groups, denoted GFP-low from sham animals and GFP-high from bleomycin-treated mice, and compared transcript levels by qRT-PCR. This analysis confirmed elevated Runx1 expression in the GFP-high population unique to the injured lung, along with elevated transcript levels for several Runx1 target genes (Fig. 3h). Collectively, these data support a role for Runx1 in promoting high collagen expression in the lung mesenchyme following injury.

### Fibrogenic memory in the Col1a2-lineage

Prompted by the residual Runx1 immunostaining we observed at 56 dpi in Col1a2-lineage cells (Fig. 3e), we next analyzed ATACseq and RNAseq datasets to determine whether any residual “memory” of fibrogenic activation persisted in the Col1a2-lineage after eight weeks, a time at which collagen levels were resolved in the lung and transcriptional changes were overwhelmingly reversed (Fig. 1e and S2). ATACseq analysis identified 908 loci that exhibited increased accessibility and 602 loci that exhibited decreased accessibility at 56 dpi relative to sham controls (Fig 4a,b). Annotating these chromatin accessibility changes to the nearest TSS revealed these changes occur distal (e.g. intergenic, intron regions) relative to TSS/promoter regions (Fig. S3a). These changes in accessibility largely represented maintenance of changes first observed at 14 dpi (Fig. S3b), demonstrating the persistence of a subset of epigenomic changes in the Col1a2-lineage following overt injury resolution. Interestingly, de novo motif enrichment analysis of differentially accessible chromatin at 56 dpi revealed Runx1 as the lead candidate regulator of persistently increased chromatin accessibility in the lung resident Col1a2-lineage at 56 dpi (Fig. 4c). Relative to the ATACseq analyses, RNAseq analyses revealed far fewer differences in transcript levels at 56 dpi; 27 genes exhibited persistently increased transcripts and 20 genes exhibited persistently decreased transcript levels (Fig. 4d). Runx1 transcript levels were not themselves significantly elevated at 56 dpi compared to sham (Fig. 4e), but increased expression of previously identified Runx1 target genes (Fig. 3) was observed (e.g. Cthrc1, Tnc). While Runx1 occupancy was previously associated with all of these loci (Fig. 4f) only Tnc and Col28a1 were observed to have persistently differential chromatin accessibility at 56 dpi (Fig 4f). Finally, we annotated these persistent chromatin accessibility changes to the nearest TSS and performed unbiased pathway analysis. The persistent epigenetic changes were in closest proximity to genes involved in pro-fibrotic gene programs, such as focal adhesions, proteoglycan production, and regulation of actin cytoskeleton (Fig. S3c). Collectively, these observations demonstrate that lung resident Col1a2-lineage retains a memory of prior fibrogenic activation. The precise genomic location of the epigenetic and transcriptional alterations at 56 dpi suggests that following injury resolution, the previously labeled lung mesenchyme exist in a “poised” state, for future, and potentially more deleterious, activation.

**Figure 4:**
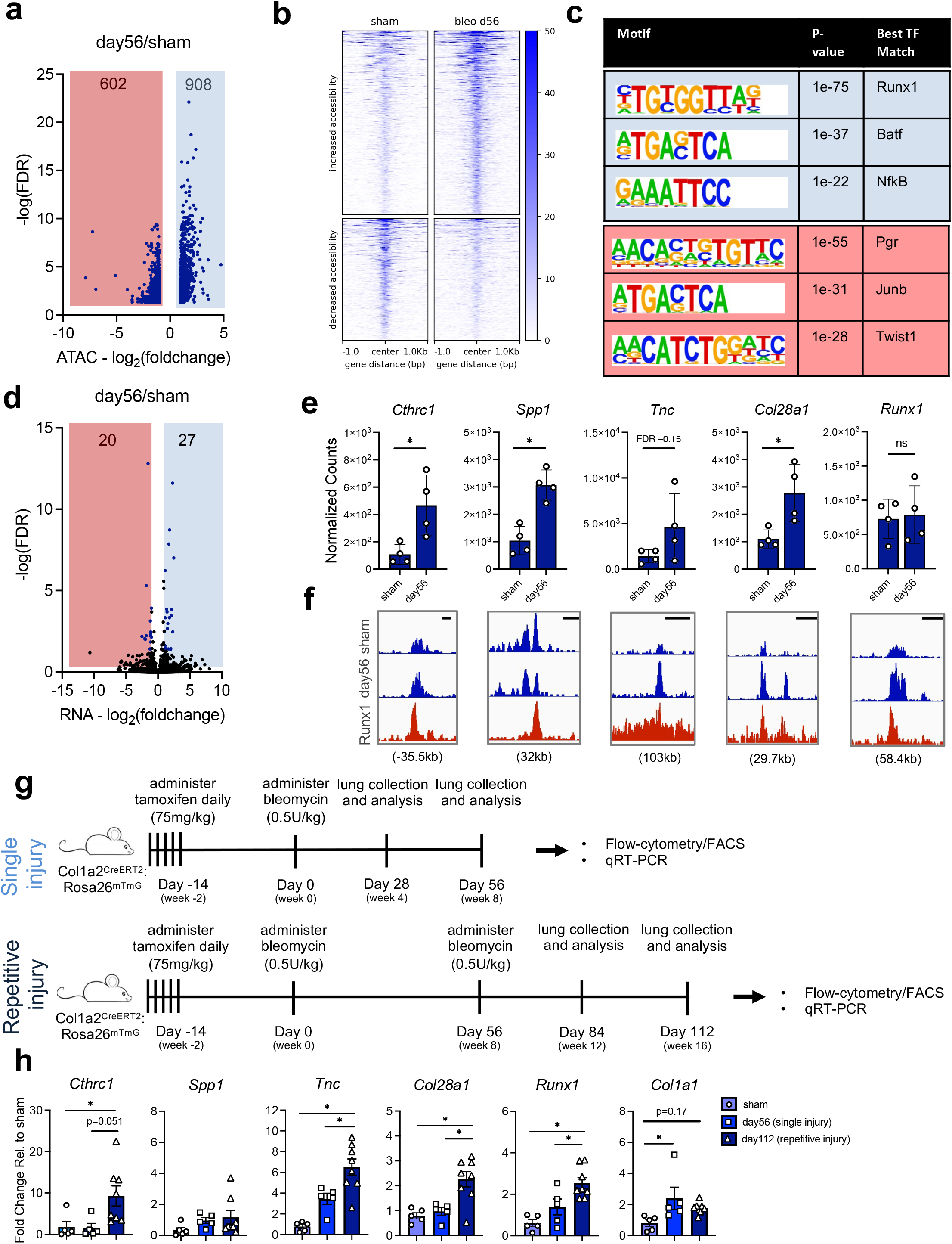
Fibrogenic memory in the Col1a2-lineage mesenchyme following single and repetitive lung injury. (a) Volcano plot of differential accessibility sites of the Col1a2-lineage at 56 dpi compared to sham (n=4 biological replicates/condition). Blue dots represent genes which met differential expression criteria (FDR ≤ 0.05 and log2(foldchange) ≤ −1 or ≥ 1). (b) Heatmap and intensity profiles of differential chromatin accessibility regions at 56 dpi compared to sham. Heatmap scale represents relative reads per genomic region. (c) De novo transcription factor motif enrichment analysis from the genomic loci with increased and decreased accessibility at 56 dpi compared to sham. Motifs ranked by p-value. (d) Volcano plot of differential gene expression of the Col1a2-lineage at 56 dpi compared to sham (n=4 biological replicates/condition). Blue dots represent genes which met differential expression criteria (FDR ≤ 0.05 and log2(foldchange) ≤ −1 or ≥ 1). (e) Candidate fibrosis-relevant genes which maintain residual changes in transcript levels at 56 dpi compared to sham. Data represent mean +/- SEM. Each dot represents a single biological replicate. *FDR<0.05. (f) ATAC and Runx1 ChIPseq profiles showing Runx1 occupancy near genes shown in panel (e). Scale bar represents 1kb in genomic distance. Number below occupancy profiles represents genomic distance from TSS. (g) Experimental approach to follow Col1a2-lineage following repetitive bleomycin lung injury. (h) qRT-PCR analysis of genes from panel (e) in addition to other candidate markers of mesenchymal cell activation (*Runx1* and *Col1a1*). Data represents mean +/- SEM. Each dot represents a single biological replicate. *P<0.05 evaluated by one-way ANOVA with Tukey’s correction for multiple comparison test.

To test whether persistent chromatin accessibility associated with Runx1-binding motifs leads to amplified responses in the Col1a2-lineage upon repeated injury, we administered bleomycin and harvested lungs at 28 and 56 dpi or subjected mice to a second dose of bleomycin at 56 dpi following the first dose (Fig. 4g). We then sorted Col1a2-lineage cells at 28 and 56 dpi after the second injury (84 and 112 days following first injury) and used qRT-PCR to compare expression of hallmark genes related to mesenchymal activation as well as Runx1 target genes at identical time points after single or repeated lung injury. Strikingly, we observed amplified responses to repeated injury at either 28 or 56 dpi for multiple Runx1 target genes, though not Col1a1 (Fig. 4h, S3d). Of note, we also observed a step-wise increase in Runx1 transcript levels upon re-injury (Fig. 4h), consistent with amplified Runx1 engagement upon repeated bleomycin exposure (Fig. 3d,e and 4f). Collectively, these data demonstrate that fibrogenic memory, potentially conferred by persistent Runx1 occupancy, primes the Col1a2-lineage for amplified responses to repetitive lung injury, contributing to fibrotic disease progression.

### Spontaneous activation of Runx1 in cultured fibroblasts

Given the prominent role for Runx1 identified in our studies above, we were curious why Runx1 activation has not been more studied in fibrogenic lung fibroblast activation. Long-term culture of mesenchymal cells has previously been shown to result in a long lasting activated cell state (Balestrini *et al.*, 2012; Yang *et al.*, 2014; Li *et al.*, 2017), and our own recent work has highlighted the spontaneous fibrogenic activation of lung fibroblasts upon encountering a stiff ECM (Jones *et al.*, 2021). To define the changes in chromatin accessibility and transcript levels experienced by lung mesenchyme when placed into standard cell culture conditions, we used a constitutive Col1a1-GFP reporter mouse to isolate the lung cells actively expressing Col1a1. We then cultured Col1a1-GFP+ cells on collagen I coated stiff substrates for a relatively brief 5 days and performed ATACseq and RNAseq, comparing our results to freshly sorted Col1a1-GFP+ cells (Fig. 5a). ATACseq analysis of freshly-sorted (“in vivo”) vs cultured (“in vitro”) Col1a1-GFP+ cells revealed striking differences between the two groups; we observed 19,809 sites with increased accessibility and 36,246 sites with decreased accessibility (Fig. 5c,d). Strikingly, a Runx1-binding motif was strongly enriched in loci exhibiting increased accessibility in vitro (Fig. 5e). RNAseq analyses revealed similarly striking differences with 4,115 genes exhibiting increased transcript levels in culture and 5,593 genes exhibiting decreased transcript levels (Fig. 5f). Interestingly, several of the same Runx1-associated genomic loci exhibited increased accessibility in vitro that also exhibited increased accessibility at 14 dpi in vivo, including those in proximity to Spp1, Tnc, and Runx1 (Fig. 5g). Transcripts for these genes also exhibited increased levels in vitro compared to fresh sorted, and at 14 dpi compared to sham (Fig. 5g, S3). Together these results are consistent with the spontaneous activation of Runx1 in cultured lung fibroblasts, an effect that may have previously masked the role that Runx1 plays in the reversible conversion of fibroblasts between quiescent and activated states in response to injury and repair.

**Figure 5:**
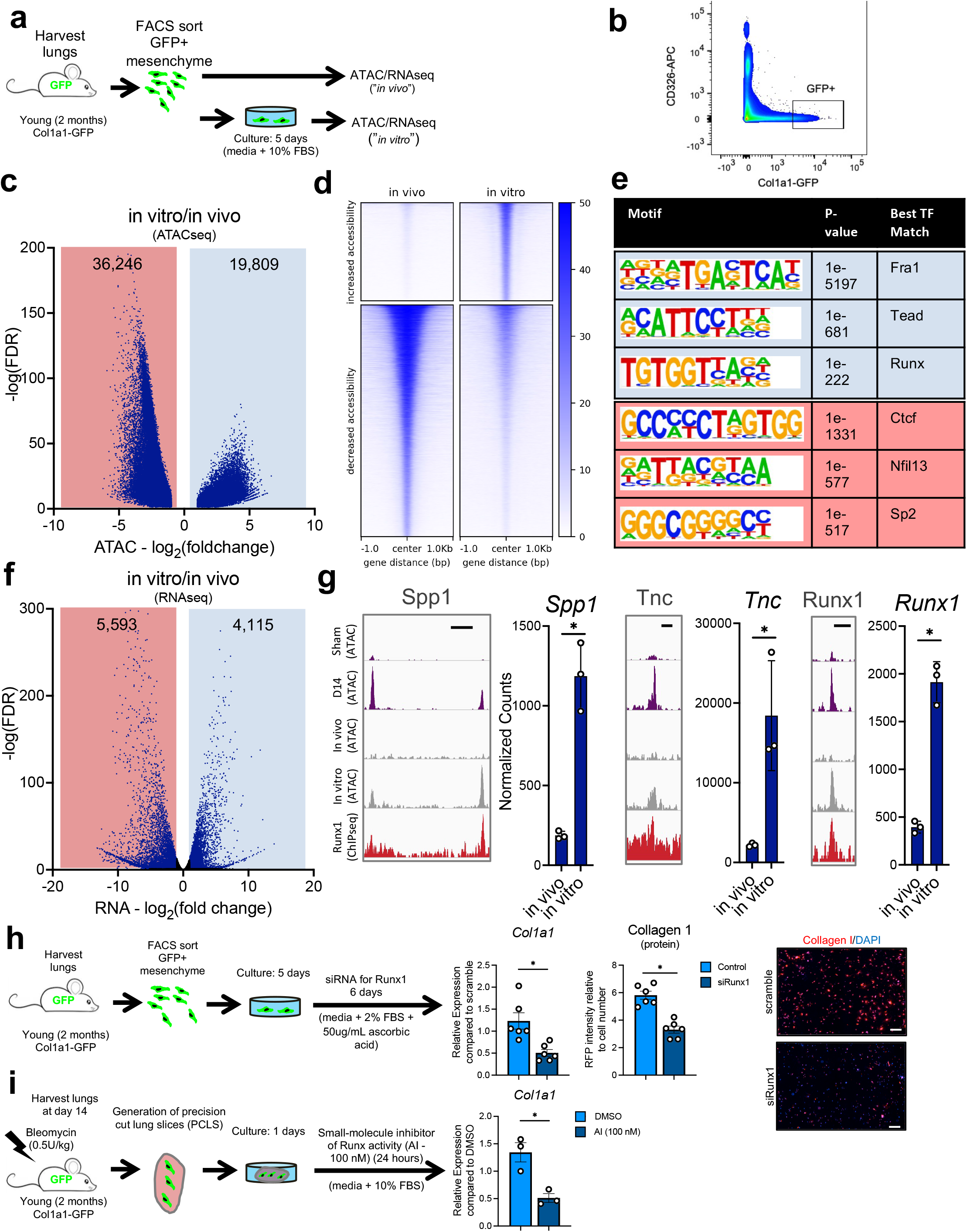
Spontaneous activation of Runx1 signaling in culture. (a) Experimental schematic showing multi-modal approach to identify mechanisms of Col1a1-GFP+ lung mesenchyme activation state in vitro compared to in vivo. (b) Representative flow-cytometry plot showing gating strategy to isolate Col1a1-GFP+ lung mesenchyme. (c) Volcano plot of differential accessibility sites of Col1a1-GFP+ lung mesenchyme in vitro compared to in vivo (n=3 biological replicates/group). Blue dots represent genes which met differential expression criteria (FDR ≤ 0.05 and log2(foldchange) ≤ −1 or ≥ 1). (d) Heatmap and intensity profiles of differential chromatin accessibility regions in vitro compared to in vivo. Heatmap scale represents relative reads per genomic region. (e) De novo transcription factor motif enrichment analysis from the genomic loci with increased and decreased accessibility in vitro compared to in vivo. Motifs ranked by p-value. (f) Volcano plot of differential gene expression of Col1a1-GFP+ lung mesenchyme in vitro compared to in vivo (n=3 biological replicates/condition). Blue dots represent genes which met differential expression criteria (FDR ≤ 0.05 and log2(foldchange) ≤ −1 or ≥ 1). (g) Representative chromatin accessibility profiles from in vitro vs in vivo differential ATAC regions and sham and 14 dpi differential ATAC regions along with Runx1 co-occupancy. Scale represents 1kb of genomic distance. RNA expression profiles are shown for the same genes. Each data point represents a single biological replicate. Data represented as mean +/- SEM. (h) Experimental approach to target Runx1 in cultured Col1a1-GFP+ lung mesenchyme using siRNA. Each dot represents a single biological replicate. Data represented as mean +/- SEM. *P <0.05, evaluated by unpaired t-test. Scale bar in immunofluorescence images represents 100 μm. (i) Experimental approach to target Runx1 by small molecule inhibitor in precision-cut lung slices prepared from Col1a1-GFP mice at 14 dpi. Each dot represents a data obtained from a single biological replicate. Data represented as mean +/- SEM. *P <0.05, evaluated by unpaired t-test.

### Runx1 inhibition reduces fibrogenic activation in vitro and ex vivo

To test whether Runx1 inhibition can attenuate fibrogenic activation of lung mesenchyme, we used both an RNA-interference (siRNA) approach and a small-molecule inhibitor of Runx1 activity (Illendula *et al.*, 2016) to block Runx1 function. Using cultured Col1a1-GFP+ cells from the mouse lung, we administered siRNA targeting Runx1 for 6 days (Fig. 5h) and measured both expression of Col1a1 and Collagen I protein. We observed that silencing of Runx1 in cultured Col1a1-GFP+ mesenchyme significantly reduced Col1a1 transcript levels and Collagen I protein (Fig. 5h).

To study the role of Runx1 in a more “in vivo”-like micro-environment, we generated precision-cut lung slices (PCLS) from Col1a1-GFP mice at 14 days following bleomycin injury. We then cultured the PCLS samples for 24 hours in the presence of a small-molecule inhibitor of Runx1 and measured Col1a1 transcript levels. Consistent with the siRNA approach, we observed that Runx1 inhibition in an “in vivo”-like setting attenuated Col1a1 expression. Collectively, these different approaches to targeting Runx1 confirm the essential role of Runx1 in regulating the fibrogenic behavior of the lung mesenchyme, consistent with prior studies linking Runx1 signaling to scar deposition, ECM remodeling, and tissue fibrosis (Koth *et al.*, 2020; Lin *et al.*, 2020; O’Hare *et al.*, 2021).

### Runx1 marks fibrogenic subsets in mouse and human lung tissue

Our results above largely relied on population level analyses in mice, leaving open the questions of whether Runx1 expression is widespread or concentrated within fibroblast subpopulations, and whether similar effects are observed in human fibrotic lung tissue. Recent single-cell RNA sequencing work using cell harvest methods optimized for the isolation of matrix-embedded mesenchyme from fibrotic tissues has demonstrated the emergence of unique subpopulations of activated mesenchyme during lung fibrogenesis (Tsukui *et al.*, 2020). To determine the distribution of Runx1 expressing cells within the lung, we downloaded publicly available scRNAseq data generated from FACS isolated Col1a1-GFP+ lung mesenchyme 14 dpi using the bleomycin model (GSE132771). Using the Seurat pipeline, we performed clustering analysis and identified 18 populations within this dataset (Fig. 6a). We next compared the expression distribution of Cthrc1, a marker of activated lung mesenchyme (Tsukui *et al.*, 2020), and Runx1 and found strong overlap of Cthrc1 and Runx1 expression in cluster 11 (Fig. 6b). Excluding immune, endothelial, and epithelial cells (Fig. S4a), we next performed differential expression analysis to identify which genes are uniquely expressed in cluster 11 compared to the other mesenchymal clusters. Consistent with prior work (Tsukui *et al.*, 2020), Cthrc1 was the top gene identified within the cluster (Fig. 6c). Remarkably, Runx1 was the top transcriptional regulator uniquely expressed in this pathological mesenchymal subpopulation (Fig. 6c). Investigation of the contribution of both “bleo” and “sham” treatment groups to the cell number of each cluster revealed cluster 11 was almost exclusively enriched for cells from bleomycin-injured mice (99.7%) (Fig. 6b). Consistent with our previous results, cluster 11 was also highly enriched for expression of other Runx1 target genes, such as Col1a1, Spp1, and Tnc (Fig. 6e).

**Figure 6:**
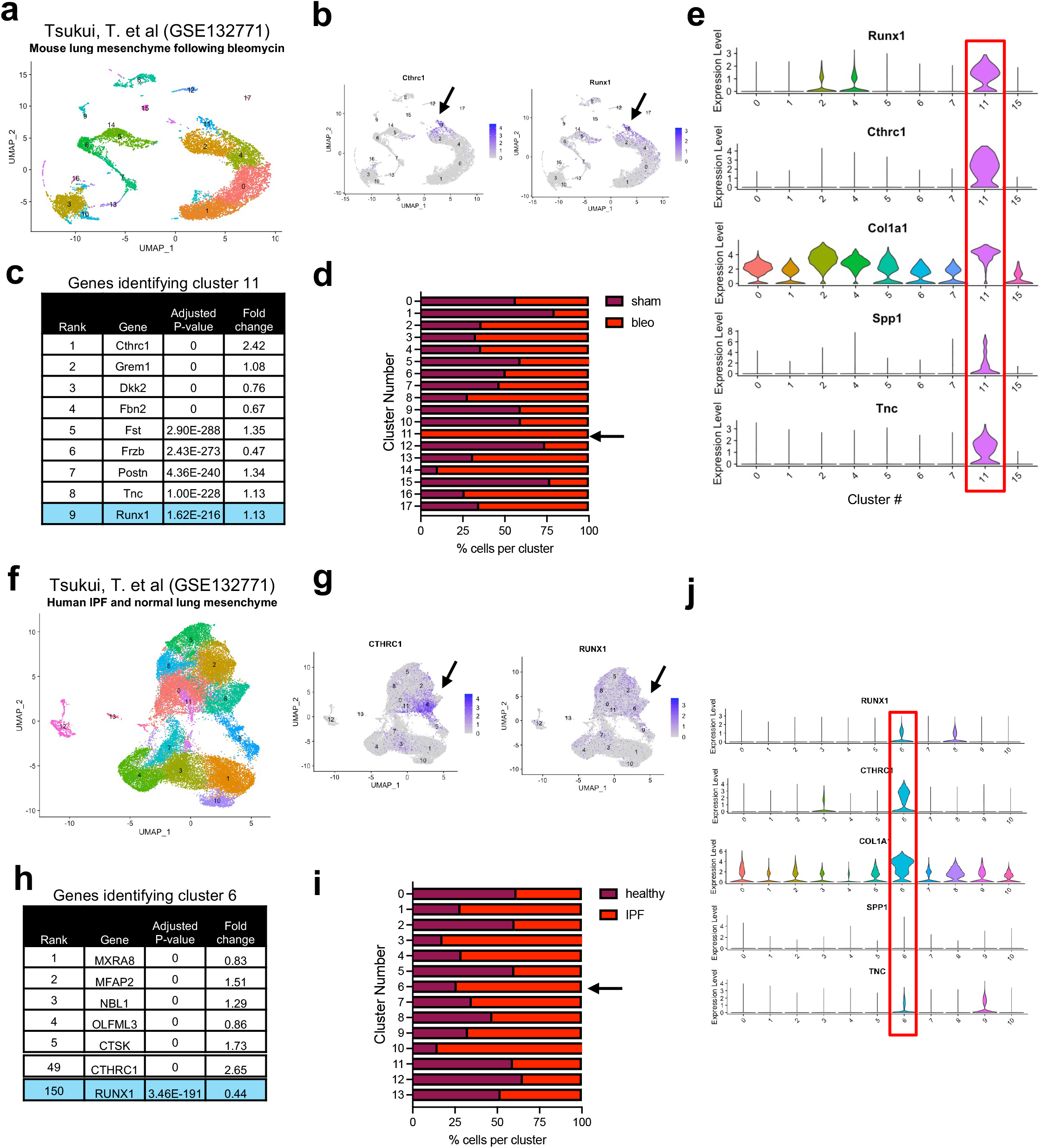
Runx1 signature in fibrotic mouse and human mesenchymal subpopulations. (a) Uniform manifold approximation and projection (UMAP) plot of all FACS isolated GFP+ cells from Col1a1-GFP mice at day 14 following bleomycin administration and sham. Data downloaded from GSE132771. (b) Gene expression profiles of Cthrc1 and Runx1. (c) Differential gene expression analysis showing genes most uniquely expressed in cluster 11 compared to other mesenchymal cell clusters. Genes ranked by adjusted p-value. (d) Contribution of treatment type (sham and bleomycin) to each cluster in panel (a). (e) Violin plots showing Runx1 target genes in each mesenchymal cell cluster. Clusters containing epithelial, endothelial, and immune cell populations were excluded. (f) Uniform manifold approximation and projection (UMAP) plot of all FACS isolated lineage-negative cells (CD45-, CD31-, and CD326-) from digested healthy and IPF human lungs. Data downloaded from GSE132771. (g) Gene expression profiles of CTHRC1 and RUNX1. (h) Differential gene expression analysis showing genes most uniquely expressed in cluster 6 compared to other mesenchymal cell clusters. Genes ranked by adjusted p-value. (i) Contribution of treatment type (healthy and IPF) to each cluster in panel (f). (j) Violin plots showing Runx1 target genes in each mesenchymal cell cluster. Clusters containing epithelial, endothelial, and immune cell populations were excluded.

To investigate Runx1 expression in the mesenchyme from IPF lung tissue, we analyzed publicly available scRNAseq from IPF and healthy human lung tissue (n=3/group) (GSE132771). Clustering analysis revealed 14 subpopulations (Fig. 6f). Expression comparison of CTHRC1 and RUNX1 again revealed an overlap of expression of these two genes within the same cluster (cluster 6) (Fig. 6g). Differential gene expression analysis revealed a significant statistical enrichment of RUNX1 in this cluster (Fig. 6h) which was composed mostly of IPF derived mesenchyme (74%) (Fig. 6i). While we did not observe strong enrichment of SPP1 in cluster 6, we did observe increased abundance of other RUNX1 target genes, such as COL1A1 and TNC (Fig. 6j). Collectively, these results confirm enrichment of RUNX1 expression in the fibrogenic subsets of mesenchyme that emerge and persist in the fibrotic mouse and human lung.

## Discussion

Our study reveals for the first time the capacity of the lung mesenchyme to retain memory of prior injury, resulting in amplified subsequent responses driven by epigenetic memory acquired during transient fibrogenesis. We show lineage-traced lung mesenchyme possess remarkable epigenomic and transcriptomic plasticity during fibrogenic activation and repair. However, following injury resolution, the lineage-labeled mesenchyme is unable to completely de-activate and thus exists in a “primed” or “poised” state. Repetitive injury experiments demonstrate a stepwise inverse relationship between number of severe injuries and the ability of the mesenchyme to de-activate. Finally, we identify Runx1 as a driver of fibrogenic memory, both in vivo and in vitro.

While the capacity of cells in other organs to retain memory of previous insults has been previously established (Kisseleva *et al.*, 2012; Naik *et al.*, 2017; Gonzales *et al.*, 2021; Larsen *et al.*, 2021), whether the lung mesenchyme possesses the capacity to retain memory has remained relatively unexplored. Our findings that the lung mesenchyme retains fibrotic-relevant memory following fibrogenic injury resolution demonstrates a new and unappreciated role of these cells in the amplified response to repetitive injury. This memory capacity of the lung mesenchyme could provide the alveolus a mechanism to more rapidly repair and regenerate following future tissue insult. However, a possible consequence of this mechanism is an increased potential for tissue fibrosis, or persistent mesenchymal cell activation, as exemplified by the amplified responses we observer after repetitive-bleomycin injury (Fig. 4g,h). It is worth noting that these bulk analyses are unable to define mesenchymal subpopulation-specific contributions to our findings. Thus, we are unable to discern whether persistent memory following resolution is dominated by a subset of mesenchymal cells, or widely shared across the broader Col1a2+ lineage spectrum. Furthermore, recent work has also highlighted other cell compartments in the lung that are persistently altered following injury resolution, some of which may provide signals to maintain fibrogenic memory in the mesenchyme (Misharin *et al.*, 2017). Given that the collagen I expressing mesenchyme is quite diverse (Tsukui *et al.*, 2020) and that bleomycin induced lung injury occurs in a regional/zonal fashion, future efforts combining multi-modal analyses within a spatial setting will be required to unambiguously address these questions. We do note that residual Runx1 immunostaining appeared to persist in the nuclei of Col1a2-lineage cells even in normal appearing regions of the lung following bleomycin injury (Fig. 3e), consistent with a cell-autonomous memory of prior fibrogenic activation.

Our finding that Runx1 is a potent regulator of mesenchymal fibrogenic function is consistent with prior reports of Runx1 regulating scar deposition and tissue fibrosis (Koth *et al.*, 2020; Lin *et al.*, 2020; O’Hare *et al.*, 2021), including a recent report demonstrating Runx1 regulating lung mesenchymal cell activation in vitro (Dubey *et al.*, 2022). Our multi-modal analyses revealed Runx1 as both a transcriptional and epigenetic regulator of Col1a2+ cell activation suggesting its role as a pioneer transcription factor (Zaret and Carroll, 2011; Hass *et al.*, 2021) in the lung Col1a2+ cell lineage. Many upstream cytokine and inflammatory signaling pathways converge on Runx1, including TGFβ and TNFα (Ito and Miyazono, 2003; Whitmore *et al.*, 2021), implicating Runx1 as an integrator of injury and fibrotic responses in the lung. How the majority of alterations in Runx1-associated chromatin accessibility changes are reversed during fibrosis resolution, and what specifies those that remain, also represents a question for further investigation. A particularly novel observation suggested by our results is the potential role of Runx1 in instilling and maintaining fibrogenic memory in the lung mesenchymal compartment following injury resolution. Interestingly, transient expression and binding of Runx1 is sufficient to induce long-term alterations in transcription (Hoogenkamp *et al.*, 2009) and other work has highlighted the interaction of Runx1 with epigenetic modifying enzymes (Reed-Inderbitzin *et al.*, 2006; Suzuki *et al.*, 2017) also implicated in fibrotic lung diseases (Huang *et al.*, 2013, 2014; Ligresti *et al.*, 2019). However, Runx1 also promotes its own expression, potentially generating a self-sustaining mechanism for maintenance of memory. Thus, discerning the extent to which Runx1 actively maintains mesenchymal memory, or whether memory persists after prior transient Runx1 activity, is open to further investigation. Answering the question of how Runx1 instills and maintains fibrogenic memory in the mesenchyme has potentially important clinical implications as we seek to therapeutically target mesenchymal memory in tissue fibrosis.

### Limitations of study

There are limitations to these experiments. With the data presented here, we cannot assess how much of the transient elevation and reversal of gene expression and chromatin accessibility is a function of activation of a subset of the Col1a2+ lineage, followed by their apoptosis versus cell epigenetic/transcriptional plasticity. It is likely a certain percentage of the activated Col1a2+ lineage undergo apoptosis following fibrogenic activation, as previous work has identified an important role for mesenchymal apoptosis in fibrosis resolution (Elizabeth F. Redente *et al.*, 2021). While our data cannot delineate the contributions of apoptosis, our repetitive injury experiments demonstrate mesenchymal cell plasticity does play a role in long-term lung tissue responses to fibrogenic injury. Because of the length of these studies, we have not studied timepoints beyond day 56 following single or repetitive injury. The durability of memory beyond the timepoints studied here remains to be determined. Our experiments focused exclusively on Col1a2-lineage labelled prior to injury, thus we cannot address whether other cells are fibrogenically activated and themselves also contribute to memory of prior activation. Moreover, while we have focused on Runx1, our analyses implicate a number of transcriptional regulators beyond Runx1 that may confer aspects of memory within the mesenchyme. Future work will be needed to build on these initial observations.

## Acknowledgements

The authors would like to thank the staff at the Mayo Clinic flow-cytometry core and microscopy core for their assistance and expertise. Funding support was provided by the National Institutes of Health grants HL105355 (to D.L.J. and P.A.L), HL159704 (to P.A.L.), HL158018 (to P.A.L.), HL153026 (to Q.T.), DK58185 and DK126827 (to T.O.), HL142596 (to G.L.), HL092961 and HL152967 (to D.J.T.). Additional funding for this work includes the Boehringer Ingelheim Discovery Award in Interstitial Lung Disease (A.J.H), ALA Catalyst Award (A.J.H.), and the Pulmonary Fibrosis Foundation Scholars Award (A.J.H.)

## Author Contributions

D.L.J., J.A.M., A.J.H, G.L., and D.J.T. designed the experiments. D.L.J., N.C., J.A.M., K.M.C., A.J.H, P.A.L., and Q.T. performed the experiments. D.L.J., N.C., K.M.C., A.J.H, and D.J.T analyzed acquired data. D.L.J. and D.J.T. wrote the first draft of the manuscript. All authors contributed to editing the manuscript.

## Methods

### Lineage tracing

All mouse experiments were carried out under a protocol approved by the Mayo Clinic Institutional Animal Care and Use Committee. The following mouse lines were used: Col1a2-CreERT (B6.Cg-Tg(Col1a2-cre/ERT,-ALPP)7Cpd/2J, Jax #029567), Rosa26-mTmG mice (B6.129(Cg)-Gt(ROSA)26Sortm4(ACTB-tdTomato,-EGFP)Luo/J, Jax #007676), and Col1a1-GFP,generated as previously described (Yata et al., 2003) and kindly provided by Dr. Derek Radisky (Mayo Clinic, Jacksonville, FL). Lineage-tracing was induced by tamoxifen injection (5 injections of 75mg tamoxifen/kg body weight, daily). Mice had access to food and water ad libitum and were on a 12h/12h light/dark cycle.

### Bleomycin-induced lung injury

Two weeks following the last injection of tamoxifen, Col1a2-CreERT2:mTmG mice were administered bleomycin (0.5U/kg) intratracheally as previously described(Caporarello *et al.*, 2019, 2020). Lungs were collected at indicated timepoints and harvested as described below. For repetitive bleomycin injury experiments, mice were administered bleomycin as described above. 56 days (8 weeks) following the first dose, mice were again administered with bleomycin (0.5 U/kg) intratracheally. Lungs were collected at indicated timepoints.

### Lung harvest and single cell suspension

At the indicated timepoints, mice were administered ketamine/xylazine solution (100 mg/kg and 10 mg/kg, respectively) injected intraperitoneally. The left ventricle of the heart was perfused with ice-cold PBS (Thermo Fisher Scientific) to remove blood content from the lung. Lungs were then immediately harvested and minced in 10-cm Petri dishes and then incubated in digestive solution (DMEM, 0.2 mg/ml Liberase DL, and 100 U/ml DNase I). Samples were digested at 37°C for 45 minutes. Digestive solution was inactivated with 1x DMEM (Thermo Fisher Scientific) containing 10% FBS (Thermo Fisher Scientific). Cell and tissue suspension was put through a 40-μm filter and centrifuged. Cell pellet was resuspended in red blood cell lysis buffer (BioLegend) for 90 seconds and then diluted in 3x volume of PBS. Cells were centrifuged and resuspended in 200 μl FACS buffer (1% BSA and 0.5 mM EDTA, pH 7.4 in PBS). The single-cell suspension was then incubated with anti-mouse CD45:PerCpCy5.5 (BioLegend; 1:200), anti-mouse CD326-APC (BioLegend; 1:200), and DAPI (Sigma-Aldrich; 1:1,000) antibodies for 30 minutes on ice.

### Fluorescence-activated cell sorting (FACS) and analysis

Samples were FACS sorted using a BD FACS Aria II (BD Biosciences, San Jose, CA, USA). To isolate GFP+ lung mesenchyme, the following selection strategy was used: debris exclusion (FSC-A by SSC-A), doublet exclusion (SSC-W by SSC-H and FSC-W by FSC-H), dead cell exclusion (DAPI by tdTomato), CD45+ cell exclusion (PerCP-Cy5.5 by tdTomato), EpCAM+ cell exclusion and isolation of GFP+ cells (APC by GFP). For the Col1a1-GFP cell isolation and analysis, an additional antibody for anti-mouse CD31:PE (BioLegend; 1:200) was used to exclude endothelial cells. For RNA analysis, GFP+ cells were sorted directly into RLT lysis buffer (Qiagen, Valencia, CA, USA). For ATACseq, GFP+ cells were sorted into FACS buffer and centrifuged at 500g for 10 minutes at 4°C. Cells were then resuspended in cryo-preservation medium (10% DMSO, 10% FBS, in RPMI-1640 (Thermo Fisher Scientific, Waltham, MA, USA)) and placed immediately into a −80°C freezer.

Population analysis was analyzed using FlowJo version 10.8.0(BD Biosciences, San Jose, CA, USA). GFP+ cell population were calculated as total % of the CD45 negative population.

### Hydroxyproline analysis

Collagen content in the lung was measured using a hydroxyproline assay kit (Biovision, Milpitas, CA, USA) according to the manufacturer’s instructions with modifications. Frozen lung tissue was fractured and homogenized in sterile water (10 mg of tissue per 100 μl H2O) and hydrolyzed in 12 M HCl in a pressure-tight, Teflon capped vial at 120°C for 3 hours followed by filtration through a 45 μm Spin-X Centrifuge Tube filter (Corning, Tewksbury, MA). The samples were dried in a Speed-Vac overnight, followed by incubation with 100 μl of Chloramine T reagent for 5 minutes at room temperature. Samples were then incubated with 100 μl of 4-(Dimethylamino)benzaldehyde (DMAB) for 90 minutes at 60 °C. The absorbance of oxidized hydroxyproline was measured at 560 nm. Hydroxyproline concentrations were calculated from a standard curve generated using known concentrations of trans-4-hydroxyl-L-proline. The total amount of protein isolated from the weighed tissues was determined by using a protein assay kit (Bio-Rad, Hercules, CA, USA). Hydroxyproline content data are expressed as μg of collagen per mg of total lung protein (μg/mg).

### Cell and tissue culture and PCLS generation

Primary mouse fibroblasts were cultured in DMEM (Thermo Fisher Scientific, Waltham, MA, USA) supplemented with 10% FBS (Thermo Fisher Scientific, Waltham, MA, USA) and Anti-Anti (Thermo Fisher Scientific, Waltham, MA, USA) unless otherwise stated. The Runx1 inhibitor AI-14-91 was generously provided by J.H. Bushweller, University of Virginia.

To generate precision cut lung slices (PCLS) we euthanized mice and perfused the lungs with PBS as stated above. We then inflated the left lobe with 10% gelatin in HBSS (with 45 μM CaCl2, 5 μM MgSO4, and 2.5 mM HEPES, final concentrations) and tied off the bronchi to prevent passive outflow. The left lobe was then placed in ice-cold HBSS and placed in the refrigerator for 10 minutes to allow the gelatin to solidify. The base of the lobe was cut off (to make a flat surface to attach to the vibratome stand). The cut end of the lobe was glued using cyanoacrylate to the vibratome stand. The remaining space in the vibratome stand was filled with 3% agarose in DMEM and returned to the refrigerator for 10 minutes for the agarose to gel. Using a Microtome (Precision Instruments Inc, VF-300 Compresstome), we cut 300 μm thick lung slices (settings: advance 5, oscillation 2, continuous mode). The PCLS were rinsed with fresh HBSS and then cultured for 24 hours in a mixed media of 50% EMEM containing 10% FBS and 50% alveolar epithelial cell medium (AEpiCM, ScienCellTM) +/- the indicated concentration of Runx1 inhibitor (AI).

### RNA isolation, qRT-PCR, RNAseq, and analysis

RNA was isolated using the RNeasy Micro Kit (Qiagen, Valencia, CA, USA) according to manufacturer’s protocol. RNA concentration was quantified using a Nanodrop spectrophotometer. cDNA was synthesized using the SuperScript VILO kit (Thermo Fisher Scientific, Waltham, MA, USA). qRT-PCR was done using FastStart Essential DNA Green Master (Roche Diagnostics, Mannheim, Germany) and analyzed using a LightCycler 96 (Roche Diagnostics, Mannheim, Germany). Primers sequences used in this study are listed in Table S1.

RNA quality was determined using the Fragment Analyzer (Agilent, Santa Clara, California, USA). RNA samples that had RQN values ≥ 8 were approved for library prep and sequencing. For the Col1a2-lineage tracing bleomycin experiment, full length double-stranded cDNA was prepared from 1 ng of total RNA according to the manufacturer’s instructions for the SMART-Seq v4 Ultra Low Input RNA Kit® (Clontech). Quantity and quality of the amplified full length double-stranded cDNA were assessed using both Qubit (Invitrogen, Carlsbad, CA) and the Bioanalyzer. Samples were diluted and 150 pg input cDNA was used with the Nextera XT DNA Library Preparation Kit (Illumina, San Diego, CA) to generate indexed libraries. Concentration and size distribution of the final libraries were determined on an Agilent Bioanalyzer DNA 1000 chip and Qubit dsDNA assay. Libraries were sequenced at six samples per lane, following Illumina’s standard protocol using the Illumina cBot and HiSeq 3000/4000 PE Cluster Kit. The flow cells were sequenced as 100 X 2 paired end reads on an Illumina HiSeq 4000 using HiSeq 3000/4000 sequencing kit and HD 3.4.0.38 collection software. Base-calling was performed using Illumina’s RTA version 2.7.7. For the in vitro vs in vivo RNAseq experiment, RNA quality was determined using the BioAnalyzer (Agilent, Santa Clara, California, USA). RNA samples that had RQN values > 6 were approved for library prep and sequencing. RNA libraries were prepared using approximately 100 ng of total RNA according to the manufacturer’s instructions for the TruSeq Stranded mRNA Sample Prep Kit (Illumina, San Diego, CA), employing poly-A mRNA enrichment using oligo dT magnetic beads. The final adapter-modified cDNA fragments were enriched by 15 cycles of PCR using Illumina TruSeq PCR primers. The concentration and size distribution of the completed libraries were determined using a Fragment Analyzer (Agilent, Santa Clara, CA, USA) and Qubit fluorometry (Invitrogen, Carlsbad, CA, USA). Libraries were sequenced at up to five samples per lane, following Illumina’s standard protocol using the Illumina cBot and HiSeq 3000/4000 PE Cluster Kit. The flow cells were sequenced as 100 X 2 paired end reads on an Illumina HiSeq 4000 using the HiSeq Control Software HD 3.4.0.38 collection software. Base-calling was performed using Illumina’s RTA version 2.7.7.

Raw fastq files were aligned to the mm10 genome using Hisat2(Kim *et al.*, 2019). Sam files were converted to Bam files using Picard SortSam. Count matrices were generated using featureCounts from the Subread package (Liao, Smyth and Shi, 2013). Count matrices were then used for differential gene expression using Deseq2 (Kim *et al.*, 2019). The threshold for significant differential expression was an FDR ≤ 0.05 and a log2 fold-change of ≥ 1 or ≤ −1. Normalized read counts were obtained from Deseq2. Panther and Ingenuity Pathway Analysis (IPA) were used to conduct gene ontology and pathway analysis, respectively.

### ATACseq and analysis

FACS sorted GFP+ cells from 2 mice were combined for a single replicate for ATACseq. Four replicates were used per group (sham, 14 dpi, and 56 dpi). For sham and 14 dpi, two replicates represent male mice and two replicates represent female mice. For 56 dpi, all four replicates represent female mice. 50,000 cells were subjected to OMNI ATACseq following the published protocol (Corces *et al.*, 2017). The size of library DNA was determined from the amplified and purified library by a Fragment Analyzer (Advanced Analytical Technologies; AATI; Ankeny, IA), and the enrichment of accessible regions was determined by the fold difference between positive and negative genomic loci using real-time PCR. The following primer sequences were used: accessibility-positive control locus: AT-P7-F: 5’-GGCTTATCCGGAGCGGAAAT −3’, AT-P7-R: 5’-GGCTGGAACAGGTTGTGTTG −3’. Accessibility-negative control locus: AT-P13-F: 5’-TCCCCTTTACTGTTTTCCTCTAC-3’, AT-P13-R: 5’-GGATTGATGAGGAAACAGCCTC-3’. The libraries were sequenced as 50 X 2 paired end reads on an Illumina HiSeq 4000 (Illumina, San Diego, CA) using the HiSeq Control Software HD 3.4.0.38 collection software. Base-calling was performed using Illumina’s RTA version 2.7.7.

Paired-end reads were mapped to the mm10 genome using bowtie2 (Langmead and Salzberg, 2012) using the “--very sensitive” option. Sam files were converted to BAM and sorted by chromosomal coordinates using Picard SortSam. Reads not aligning to Chr1-20, X, or Y were removed using SAMtools (Langmead and Salzberg, 2012). PCR duplicates were removed from Bam files using Picard MarkDuplicates. Peaks were called using MACS2 with the following options “-q 0.01 --shift −37 --extsize 73 --keep-dup all”. Peaks which were observed in the blacklisted regions of the genome (Zhang *et al.*, 2008) were removed using Bedtools (Quinlan and Hall, 2010). DiffBind was used for examination of differential chromatin accessibility(Stark and Brown, 2011). The threshold for significantly differential accessibility was an FDR ≤ 0.05 and a log2 fold-change of ≥ 1 or ≤ −1. Differential accessibility sites were then annotated to their nearest transcriptional start site (TSS) using Homer (Heinz *et al.*, 2010). Motif analysis of the differential accessibility sites was performed using Homer using the following options “-size given” and “-mask”.

### Chromatin immunoprecipitation (ChIP)-sequencing analysis

Raw Runx1 ChIPseq data was downloaded from GSE 90893 (Chronis *et al.*, 2017). Fastq files were aligned to the mm10 mouse genome using Bowtie2 with default parameters. Sam files were converted to bam with Picard SortSam. Duplicates were removed with Picard MarkDuplicates. Peaks were called using MACS2 using a q-value threshold of 0.05. Runx1 peaks which were observed in the blacklisted regions of the genome were removed using Bedtools. To identify co-occupancy regions of Runx1 with ATACseq data, mergePeaks.pl was used within Homer. Peaks were considered co-occupied if peak summits were with 300 bp of each other.

### Single-cell RNAseq analysis

Previously published scRNAseq data from FACS isolated Col1a1-GFP lung mesenchyme following bleomycin challenge, along with freshly isolated healthy and IPF human lung mesenchyme, were downloaded from GSE132771 (Tsukui *et al.*, 2020). scRNAseq data were analyzed using Seurat v4.0 (Hao *et al.*, 2021). DimPlot function was used to generate UMAP plots. FeaturePlot was used to visualize gene transcript abundance across UMAP space. VlnPlot function was used to generate violin plots. FindMarkers function was used to identify genes marking a certain cell cluster. These analyses were performed on the mesenchymal cell populations; we excluded immune, epithelial, and endothelial populations using expression of Ptprc (Cd45), Epcam (Cd326), and Pecam1 (Cd31), respectively.

### Immunofluorescence staining

Lungs were perfused via left ventricle with cold PBS and inflated by intra-tracheal instillation of fresh 4% paraformaldehyde under a constant pressure of 25cmH2O. Lungs were then harvested and fixed with 4% paraformaldehyde overnight at 4°C, followed by cryopreservation with 30% sucrose until sinking, and embedded in OCT. Tissue sections (7 um) from each block were cut in a cryostat at −21°C and mounted onto Vectabond-coated slides (Vector Laboratories, Peterborough, UK). Slides were permeabilized in 0.25 % Triton X-100 (Sigma-Aldrich, St. Louis, MA, USA), blocked with 1% BSA for 1 hour and incubated overnight with an anti-RUNX1/AML rabbit primary antibody (Abcam, Cambridge, MA, USA, Cat# ab23980) diluted 1:500 in PBS with 1% BSA and 0.1% Triton, followed by fluorescence-conjugated secondary antibody (Thermo Fisher Scientific, Waltham, MA, USA, Cat# A31573, dilution 1:600, 1h at room temperature) and DAPI (Thermo Fisher Scientific, Waltham, MA, USA, Cat#62248, dilution 1:1000, 1h at room temperature) to counterstain nuclei. All images were captured using a Zeiss LSM 710 confocal microscope.

---

**Figure S1: Inducible labeling of lung Col1a2+ mesenchyme.** Lung tissue from non-tamoxifen treated and tamoxifen treated Col1a2-CreERT2:Rosa26-mTmG mice were digested and analyzed by flow cytometry.

**Figure S2: Transcriptomic analysis of the lung Col1a2-lineage at 14 and 56 dpi.** (a) Volcano plot of differential gene expression of the Col1a2-lineage at 14 dpi compared to sham (n=4 biological replicates/condition). Blue dots represent genes which met differential expression criteria (FDR ≤ 0.05 and log2(foldchange) ≤ −1 or ≥ 1). (b) Pathway analysis of genes which were upregulated at 14 dpi compared to sham (blue). (c) Pathway analysis of genes which were downregulated at 14 dpi compared to sham (red). (d) Volcano plot of differential gene expression of the Col1a2-lineage at 56 compared to 14 dpi (n=4 biological replicates/condition). Blue dots represent genes which met differential expression criteria (FDR ≤ 0.05 and log2(foldchange) ≤ −1 or ≥ 1). (e) Heatmap of representative differentially regulated genes at 14 dpi compared sham. Relative expression profiles at 56 dpi also shown. Each column represents a single biological replicate.

**Figure S3: Fibrogenic memory in the Col1a2-lineage following lung injury** (a) Distribution of genomic location of the differential accessibility chromatin sites which exhibited changes at 56 dpi compared to sham. (b) Number of differential accessibility regions at 56 dpi (compared to sham) which were also differentially accessible at 14 dpi (compared to sham). (c) Pathway analysis of genes nearest differentially accessible chromatin at 56 dpi compared to sham. (d) qRT-PCR analysis of FACS isolated Col1a2-lineage labeled lung mesenchyme following repetitive bleomycin injury. Each dot represents data obtained from a single biological replicate. Data represented as mean +/- SEM. *P <0.05, evaluated by one-way ANOVA with Tukey’s correction for multiple comparison test.

**Figure S4: Immune, endothelial, and epithelial populations in the mouse and human lung** (a) Expression pattern of markers of immune, endothelial, and epithelial cell markers (Ptprc, Pecam1, and Epcam, respectively) in mouse cells justifying exclusion of these clusters from Figure 6. (b) Expression pattern of markers of immune, endothelial, and epithelial cell markers (Ptprc, Pecam1, and Epcam, respectively) in human cells justifying exclusion of these clusters from Figure 6.

