## Supplementary figures and images for "Lung injury epigenetically primes mesenchyme for amplified activation upon re-injury"

### Figure S1

#### Figure S1

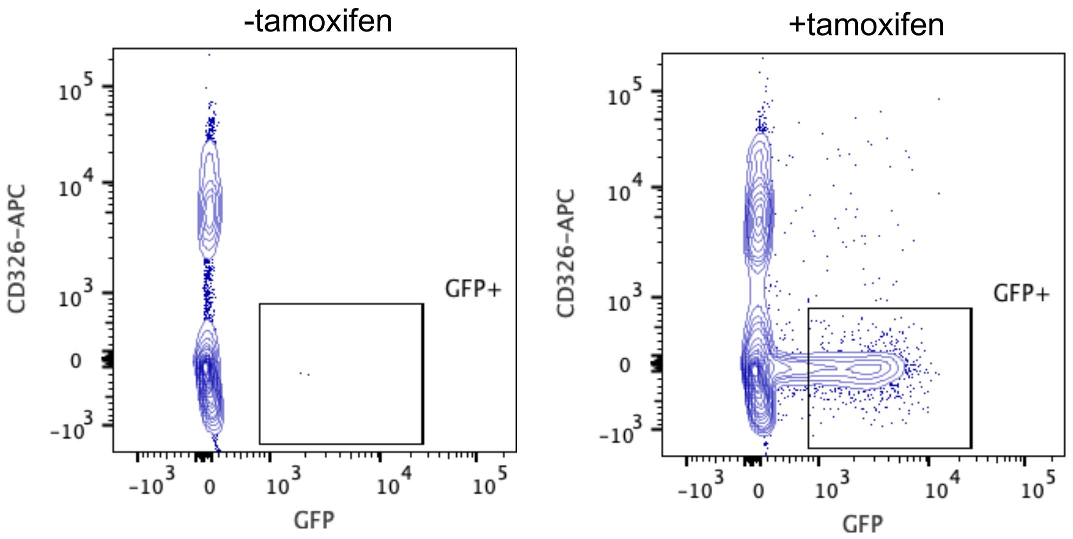

### Figure S2

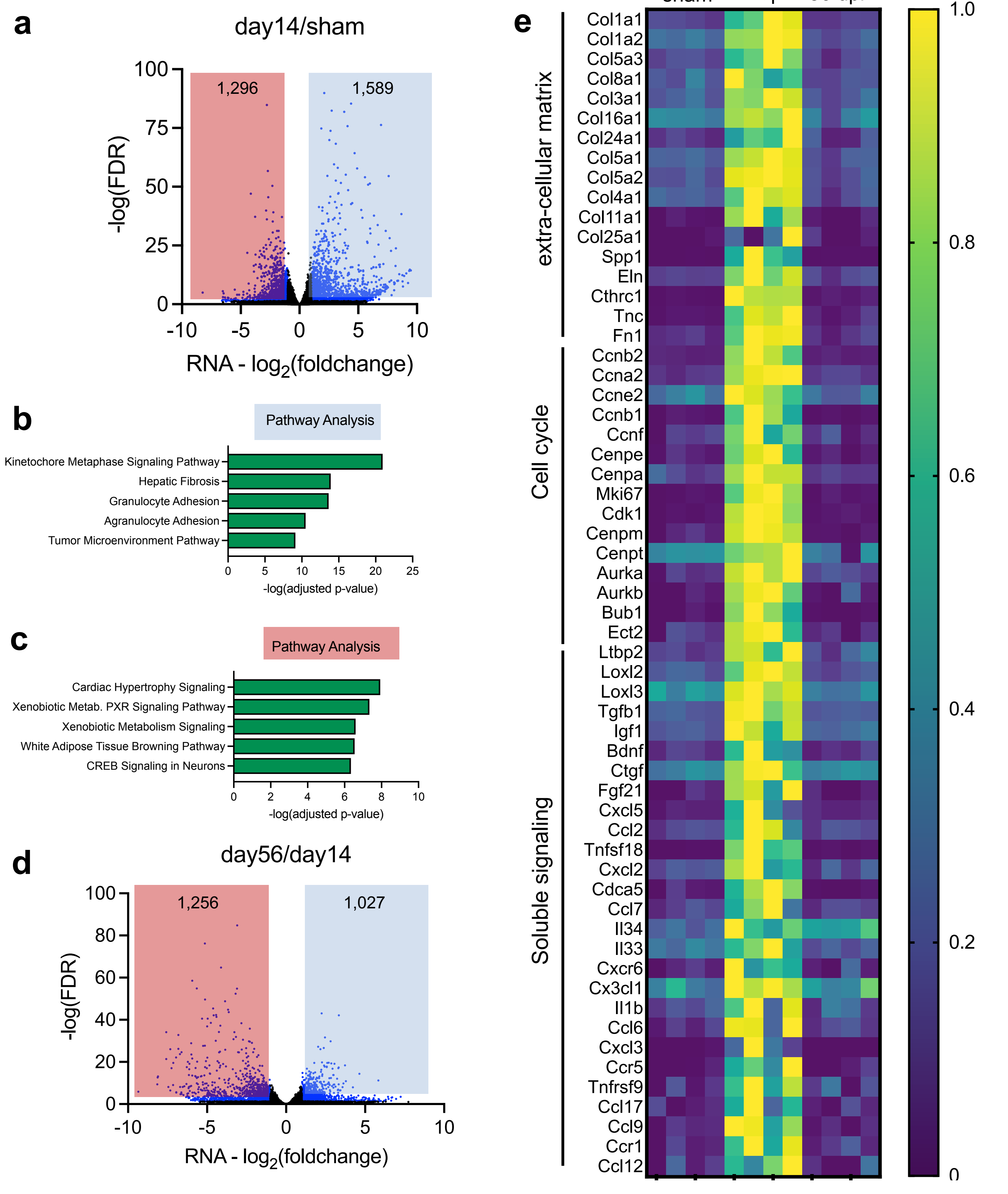

### Figure S3

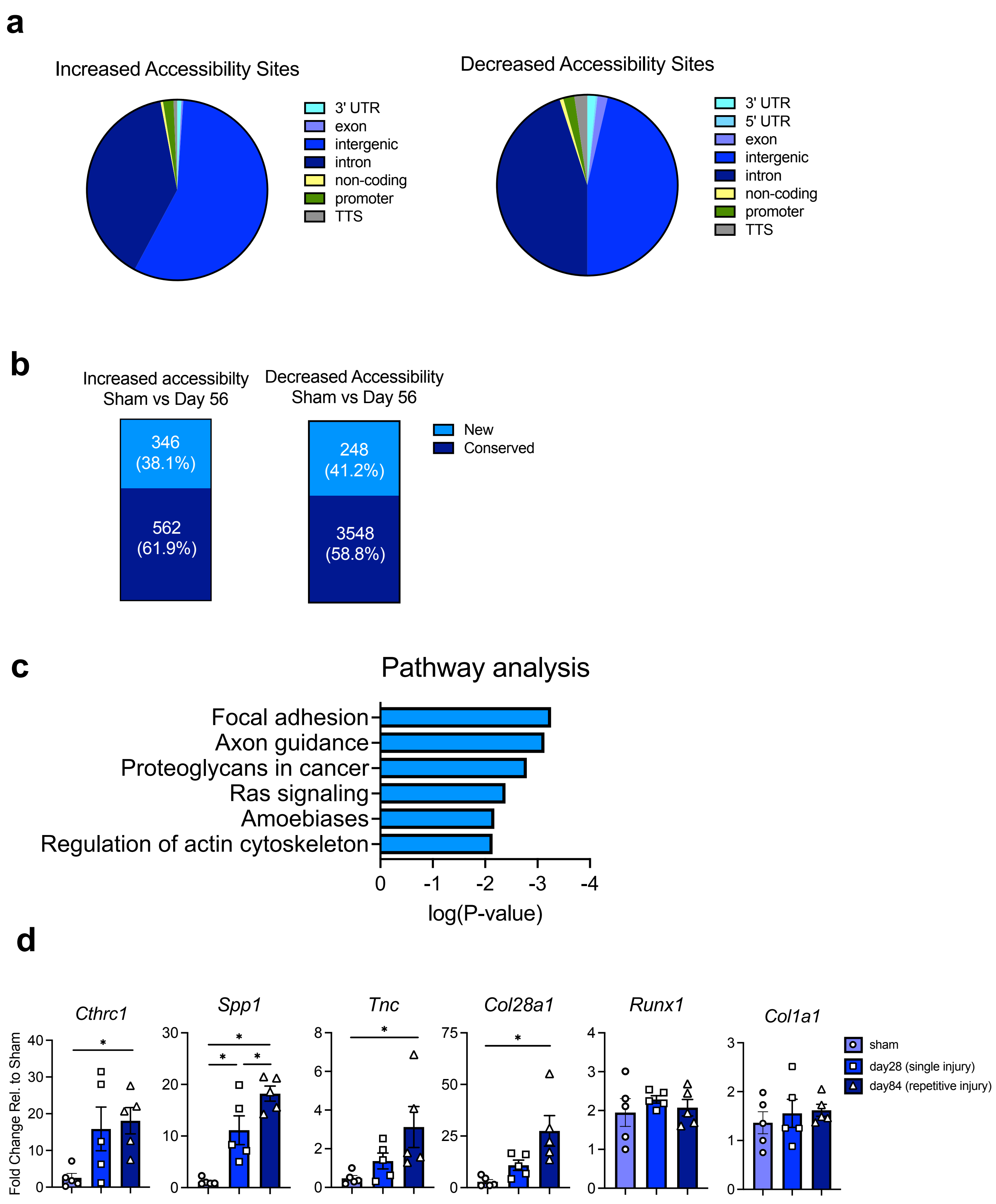

### Figure S4

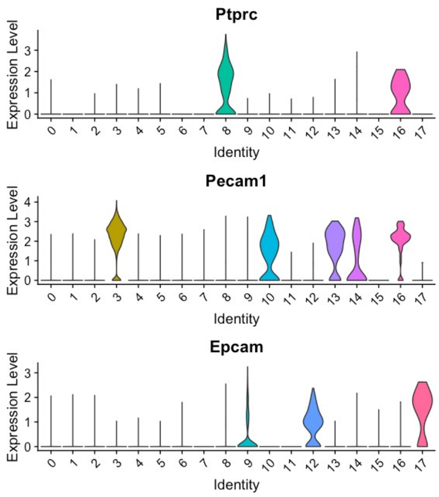

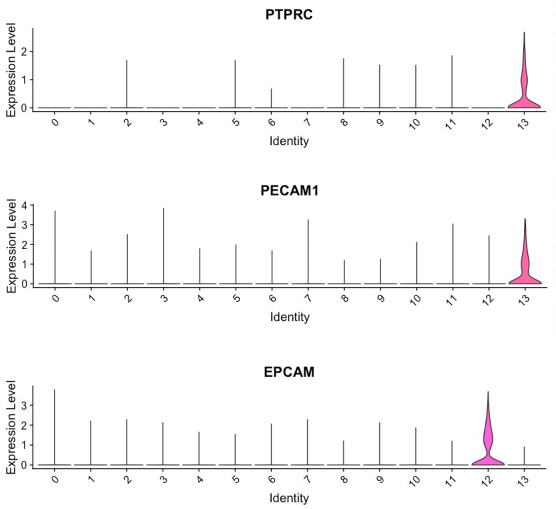
